# Allosteric modulation of β1 integrin through the hybrid domain reverses articular cartilage injury and functional impairment in a murine model of inflammatory arthritis

**DOI:** 10.64898/2026.07.09.737517

**Authors:** Rehab AlJamal-Naylor, Nicola Barton, Susan McIntyre, Daniel S. McQueen, David J. Harrison

**Author notes:** Corresponding author. (R. AlJamal-Naylor).

## Abstract

Rheumatoid arthritis is a chronic inflammatory joint disease in which progressive destruction of cartilage and bone drives long-term disability. Current disease-modifying therapies target the immune and cytokine networks that sustain synovial inflammation, but none is directed at the chondrocyte, the resident cell responsible for maintaining cartilage matrix. Chondrocyte survival and matrix homeostasis depend on β1-integrin-mediated adhesion to the extracellular matrix, and dysregulated integrin signalling has been implicated in cartilage injury. Here we test the hypothesis that allosteric modulation of β1 integrin, rather than simple adhesion blockade, is chondroprotective. Using the monoclonal antibody JB1a, which binds an epitope in the hybrid domain of β1 integrin and stabilises the receptor in a low-affinity conformation, we show that intra-articular administration produces both functional and structural amelioration of Freund’s complete adjuvant (FCA)-induced arthritis in mice. JB1a abolished the FCA-induced increase in joint diameter and hyperalgesia and markedly reduced synovial inflammation, pannus formation and cartilage erosion, with no effect on the contralateral joint and no observed adverse effects. These changes were accompanied by a reduction in chondrocyte apoptosis in vivo. In primary human articular chondrocytes, JB1a abolished interleukin-1β (IL-1β)-induced caspase 3/7 activation, reduced IL-8 secretion, and restored the sinusoidal oscillation of intracellular ATP that was otherwise abrogated by IL-1β. In contrast, the adhesion-blocking, integrin-clustering antibody 6S6 activated caspase 3/7 and amplified IL-1β-induced IL-8 secretion, indicating that the therapeutic effect is a property of the specific mode of receptor engagement rather than of adhesion blockade per se. These findings identify β1-integrin conformational state as a determinant of chondrocyte energy homeostasis and survival, and nominate allosteric β1-integrin modulation as a mechanistically distinct, chondrocyte-directed therapeutic strategy in inflammatory arthritis.

## Introduction

Rheumatoid arthritis (RA) affects approximately 0.5–1% of the population in developed countries and is characterised by synovial proliferation and subintimal infiltration of inflammatory cells [1,2]. This proliferative process is accompanied by angiogenesis and the formation of pannus, an invasive synovial tissue that is thought to contribute both to the chronicity of the inflammation and to the erosion of bone and thinning of cartilage [1]. Although the pathogenesis of RA is incompletely understood, CD4+ T-cell-mediated autoimmune responses, and in particular interferon-γ-producing T-helper-1 cells, are considered pivotal in the initiation and maintenance of autoimmune arthritis in both humans and animal models [1,2].

Current disease-modifying anti-rheumatic drug therapies slow joint destruction in only a proportion of patients, and much recent effort has focused on the specific modulation of the inflammatory response by targeting cytokines, their receptors, and downstream signalling enzymes; blockade of phosphoinositide-3-kinase-γ, for example, suppresses joint inflammation and damage in murine models of RA [3]. These approaches share a common feature: they are directed at the immune and stromal drivers of inflammation rather than at the chondrocyte, the resident cartilage cell whose loss underlies irreversible joint failure. To date there has been essentially no therapy directed at protecting the chondrocyte itself.

Articular cartilage is an avascular tissue devoid of nerves and lymphatics. The homeostasis of its extracellular matrix (ECM) depends on continuous, bidirectional communication between chondrocytes and their surrounding matrix through cell-surface adhesion receptors, principally the integrins [4]. Normal chondrocytes express α1β1, α5β1, αVβ5, αVβ3 and α3β1 integrins [4]. The interaction of β1 integrin with the ECM is essential during mesenchymal condensation and chondrocyte differentiation: administration of anti-β1 integrin antibodies to mouse limb mesenchymal buds inhibits the formation of cartilage nodules, and both inhibition of β1 integrin in cartilage explants and conditional deletion of the β1 integrin gene in mice reduce growth and type-X collagen deposition, producing abnormal chondrocyte shape, disorganised F-actin and increased apoptosis [5,6]. In arthritis-affected cartilage, α5β1 and αVβ3 integrins reciprocally regulate the production of inflammatory mediators, linking integrin engagement directly to the inflammatory phenotype of the chondrocyte [7].

We have previously proposed that β1 integrin is not merely an adhesion molecule but a druggable, conformationally regulated signalling hub whose set-point can be shifted pharmacologically [8,9]. Integrins are bidirectional, allosteric signalling machines whose affinity for ligand is governed by large-scale conformational rearrangements of the ectodomain [10]. We reasoned that, in injury, fragmentation of the ECM may lead to inappropriate β1 integrin activation and clustering, with consequent cytoskeletal reorganisation, mitochondrial dysfunction, altered ATP homeostasis and increased susceptibility to apoptosis; failure to clear the resulting apoptotic debris could in turn act as a pro-inflammatory stimulus. In this framework, controlled disengagement or conformational restraint of β1 integrin—rather than immunosuppression—could be of therapeutic value. Preliminary work indicated that specific inhibitory anti-β1 integrin antibodies increase ECM proteoglycan content in human chondrocytes (data not shown). Here we test this hypothesis in vivo and in vitro using JB1a, a monoclonal antibody directed against the hybrid domain of β1 integrin that acts as an allosteric modulator rather than a simple adhesion blocker.

## Materials and Methods

### Ethics statement

All animal procedures were carried out in accordance with the UK Animals (Scientific Procedures) Act 1986 and associated institutional guidelines, under the appropriate Home Office project and personal licences, and were approved by the local ethical review committee of the University of Edinburgh. Primary human articular chondrocytes were obtained from a commercial supplier and used in accordance with the supplier’s terms and applicable ethical approvals for the use of human-derived material.

### Induction of arthritis in vivo

Unilateral arthritis was induced by intra-articular administration of Freund’s complete adjuvant (FCA) into the stifle (knee) joint, adapting a previously characterised murine adjuvant-arthritis model [11,12]. Animals received either FCA (*Mycobacterium tuberculosis* in paraffin oil; Sigma) or vehicle (heavy liquid paraffin oil; controls) in a volume of 20 µl. Briefly, animals were transiently anaesthetised (3% halothane in oxygen), a small incision was made over the stifle joint to expose the patellar tendon, and FCA, vehicle or antibody (20 µl) was injected beneath the patellar tendon directly into the synovial space using a 30-gauge needle mounted on a 50-µl Hamilton syringe. Animals were treated with JB1a (mouse anti-human IgG1; Chemicon), isotype control (IgG1, MOPC21; Sigma) or vehicle at weekly intervals on days 7 and 14 after FCA. Sham-operated animals were also included.

### Assessment of arthritis

Animals were weighed and the joint diameter, measured across the joint just below the level of the patella, was recorded weekly (left and right) prior to injection on the same day, according to previously described methods [12,13]. Hyperalgesia was scored using a subjective pressure-application scale correlating an increased hyperalgesia score with reduced applied pressure, from 0 (normal; high pressure tolerated) to 3 (severely hyperalgesic; withdrawal at very low pressure). Testing involved gently compressing the joint between thumb and forefinger and determining the pressure required to elicit limb withdrawal (normal end-point) or, rarely, vocalisation.

### Histopathology

Animals were killed by CO2 asphyxiation on day 21 and the left (injected, ipsilateral) and right (control, contralateral) stifle joints were removed for histology. Tissues were fixed for 48 h in 10% neutral-buffered formalin, decalcified in 15% EDTA (pH 7) in 10% neutral-buffered formalin for 2 weeks, embedded in paraffin wax, sectioned at 8 µm, and stained with haematoxylin/eosin and with alcian blue (pH 1.0)/brilliant red. Images were digitised using Image-Pro Plus (version 5.1) and a MicroPublisher 3.3 RTV camera on a Zeiss Axioskop (5× objective for haematoxylin/eosin; 20× objective for alcian blue). The severity of synovial inflammation, joint damage, metachromasia and pannus formation was scored on an analogue scale (0 = no abnormality detected to 3 = very marked) by an assessor blinded to treatment, as previously described.

### Apoptosis measurement

Terminal deoxyribonucleotidyl-transferase-mediated dUTP nick-end labelling (TUNEL) was performed on sections using the ApopTag Red kit (Chemicon). Positively stained apoptotic nuclei were imaged using the 40× oil objective of a Zeiss LSM 510 confocal microscope (Carl Zeiss Ltd, Welwyn Garden City, UK). Images were converted to 8-bit greyscale and total cell number was determined using ImageJ (NIH); TUNEL-positive cells were counted manually by a blinded observer.

### Human articular chondrocytes in vitro

Primary normal adult human articular chondrocytes derived from the knee joint were cultured according to the supplier’s instructions. Cells were cultured for 2 weeks in 1.2% alginic acid (Keltone LV) in normal saline (3 weeks for bead experiments), polymerised with 102 mM CaCl2. On the day of experiment, cells were released from alginate with 55 mM sodium citrate and allowed to attach to plastic for 6 h before being serum-starved in DMEM/F12 containing 0.1% fetal calf serum (FCS) for 1 h. Carrier-free IL-1β (R&D Systems) was added at 10 ng/ml alone or in combination with JB1a (1 µg/ml) or the adhesion-blocking anti-β1 integrin antibody 6S6 (1 µg/ml). At 3, 6, 12 or 24 h the medium was aspirated and preserved, and caspase 3/7 activity was assayed using Caspase-Glo 3/7 (Promega) according to the manufacturer’s instructions. Secreted IL-8 was measured in preserved media by ELISA.

### ATP measurement

In separate experiments, chondrocytes were dispersed and seeded into 96-well plates as above, serum-starved in medium containing 0.1% FCS and then in glucose-free DMEM with 0.1% FCS, before treatment with (i) IL-1β (10 ng/ml) alone, (ii) IL-1β (10 ng/ml) plus JB1a (1 µg/ml), (iii) JB1a (1 µg/ml) alone, or (iv) vehicle control. From the third hour, intracellular ATP levels were recorded minute-by-minute using a bioluminescent ATP assay (PerkinElmer).

### Data analysis and statistics

Data were analysed using Microsoft Excel, GraphPad Prism and SPSS. One-way and multi-way ANOVA followed by unpaired *t*-tests were used to compare means of normally distributed groups. Where sample sizes were small or data were not normally distributed, the non-parametric Mann–Whitney *U*-test was used. Where a two-way ANOVA showed no significant effect of time, data for a given treatment were pooled across time points. Repeated measures were analysed by repeated-measures ANOVA with an appropriate post-hoc test. Medians of two or more non-parametric groups were compared using the Kruskal–Wallis test with Dunn’s multiple-comparison post-hoc analysis. A *P* value below 0.05 was considered significant, and exact *P* values are quoted where relevant. Values are presented as means or as scatter.

## Results

### JB1a reverses joint swelling and hyperalgesia in vivo

Mice injected with FCA showed a significant increase in ipsilateral joint diameter and hyperalgesia score on days 7, 14 and 21 relative to vehicle-treated or sham animals (Fig 1A, 1B). In animals that received FCA followed by intra-articular JB1a (60 µg) on days 7 and 14, the increases in ipsilateral joint diameter and hyperalgesia score were abolished by days 14 and 21. Neither the contralateral joint diameter nor the contralateral hyperalgesia score changed significantly with treatment (Fig 1C), indicating a local rather than systemic effect. No adverse effects were observed following JB1a administration.

**Fig 1.**
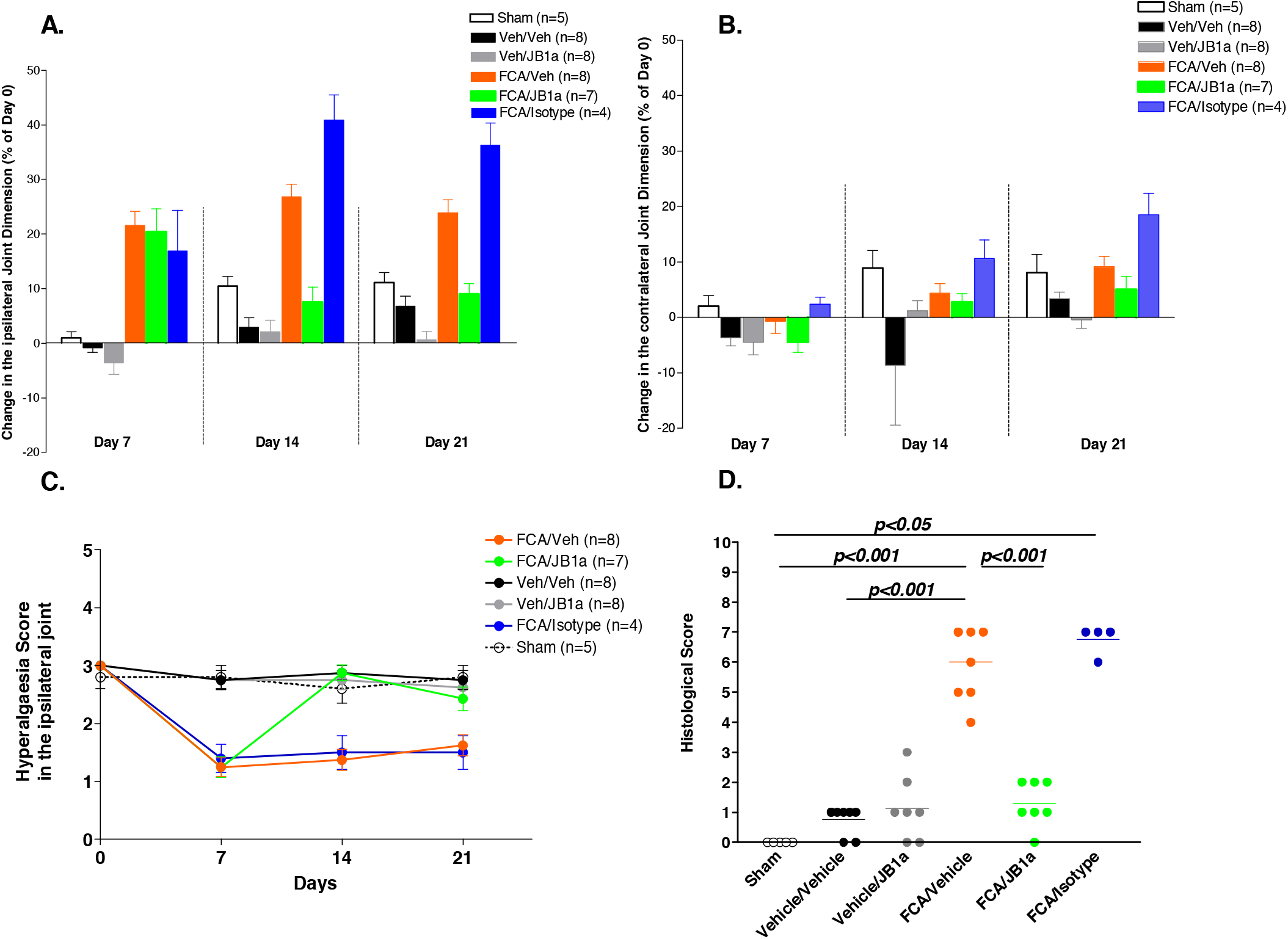
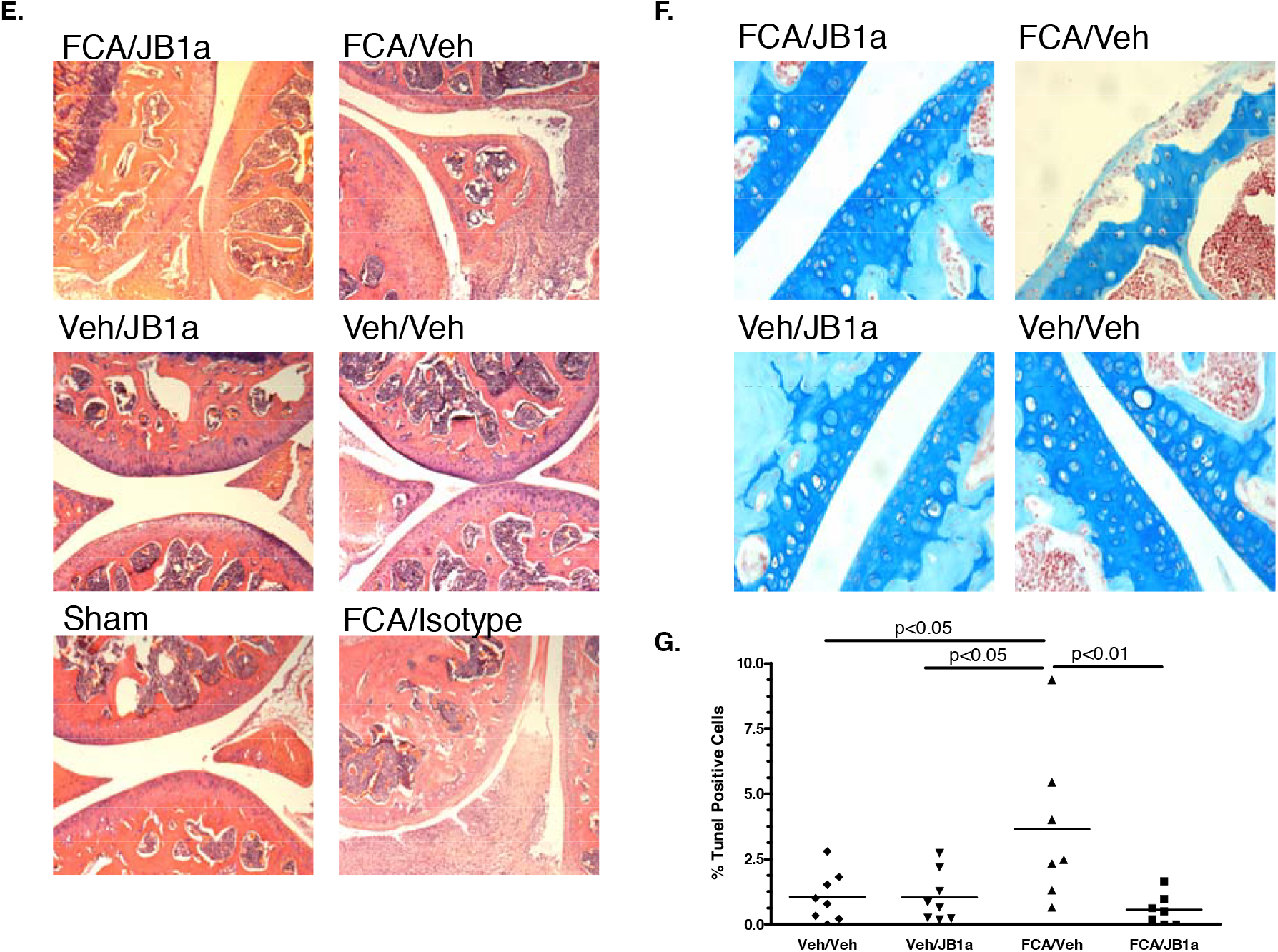
Allosteric β1-integrin modulation with JB1a in the FCA arthritis model and in IL-1β-challenged human chondrocytes. (A) Panel position reserved (see note below). (B) Intracellular ATP (relative luminescence units) recorded minute-by-minute in primary human articular chondrocytes treated with vehicle (Control), JB1a, IL-1β or JB1a + IL-1β; IL-1β dampens the oscillation, which is restored by co-treatment with JB1a. (C) IL-8 concentration (pg/ml) at 3, 6 and 12 h for Control, JB1a, IL-1β, JB1a + IL-1β, 6S6 and 6S6 + IL-1β; JB1a reduces IL-1β-induced IL-8, whereas 6S6 amplifies it. (D) Histological damage score (0–3) for Sham, Vehicle/Vehicle, Vehicle/JB1a, FCA/Vehicle, FCA/JB1a and FCA/Isotype joints, with indicated significance; FCA increases the score and JB1a returns it towards baseline. (E) Representative haematoxylin/eosin sections (FCA/JB1a, FCA/Veh, Veh/JB1a, Veh/Veh, Sham, FCA/Isotype) showing FCA-induced synovial proliferation, cellular infiltrate, pannus formation and cartilage erosion and their amelioration by JB1a. (F) Representative Alcian blue sections (FCA/JB1a, FCA/Veh, Veh/JB1a, Veh/Veh) showing preservation of matrix proteoglycan staining with JB1a. (G) Repeat of the intracellular ATP trace in panel B.

### JB1a ameliorates the histological features of arthritis

Histologically, FCA-injected joints displayed inflammation with cellular infiltrate, synovial proliferation, pannus formation and cartilage erosion (Fig 1D–1F). Joints from JB1a-treated animals showed a marked amelioration of these features, with reduced synovial thickness and cellular infiltrate, reversal of cartilage thinning and reduced pannus formation. Because JB1a abolished hyperalgesia in parallel with these structural changes, the reduction in hyperalgesia is unlikely to be explained solely by a direct effect on nociception, and is consistent with resolution of the underlying joint pathology.

### JB1a reduces chondrocyte apoptosis in the arthritic joint

We next assessed apoptosis in situ, reasoning that chondrocyte apoptosis and the failure to clear apoptotic bodies could itself propagate inflammation and autoimmunity. The background prevalence of apoptosis was low. FCA induced a significant increase in TUNEL-positive cells in the ipsilateral joint, and this increase was significantly reduced by treatment with JB1a (Fig 1G).

### JB1a abolishes IL-1β-induced chondrocyte apoptosis in vitro, whereas a clustering antibody promotes it

To interrogate the role of β1 integrin in chondrocyte injury directly, primary human articular chondrocytes were exposed to IL-1β (10 ng/ml), which produced a marked increase in caspase 3/7 activity (Fig 2A). JB1a alone had no effect on caspase 3/7 activity; however, when added together with IL-1β, JB1a abolished IL-1β-induced caspase 3/7 activation. Equivalent results were obtained in chondrocytes cultured in alginate beads for 3 weeks (data not shown). In contrast, the adhesion-blocking anti-β1 integrin antibody 6S6, which is known to induce integrin clustering and homotypic aggregation, itself activated caspase 3/7. The epitope recognised by 6S6 is discontinuous and has not been mapped. Thus, two antibodies that both block adhesion produced opposite effects on chondrocyte survival, indicating that the outcome is governed by the mode of receptor engagement rather than by adhesion blockade alone.

**Fig 2.**
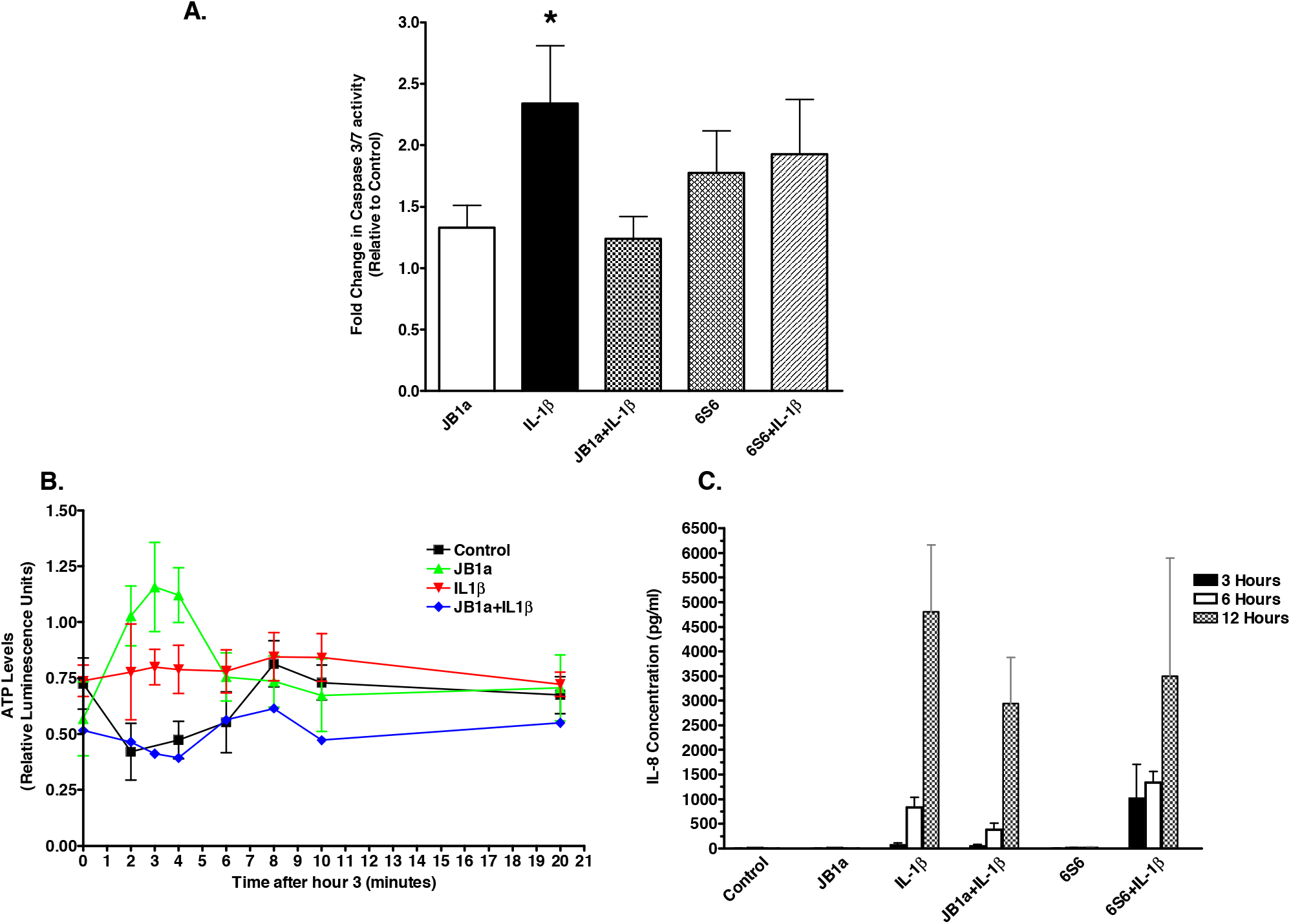
JB1a protects human articular chondrocytes from IL-1β-induced injury in vitro. (A) Panel position reserved (see note below). (B) Minute-by-minute intracellular ATP (relative luminescence units) in chondrocytes treated with vehicle (Control), JB1a, IL-1β or JB1a + IL-1β: IL-1β abrogates the sinusoidal oscillation seen in control cells, and co-treatment with JB1a restores it. (C) IL-8 concentration (pg/ml) at 3, 6 and 12 h for Control, JB1a, IL-1β, JB1a + IL-1β, 6S6 and 6S6 + IL-1β: IL-1β induces time-dependent IL-8 release that is reduced by JB1a and amplified by 6S6.

### JB1a attenuates IL-1β-induced IL-8 secretion

IL-1β stimulated the secretion of IL-8 in a time-dependent manner (Fig 2B). This response was reduced by approximately 50% in the presence of JB1a. Although 6S6 had no effect on IL-8 secretion when applied alone, it amplified IL-1β-induced IL-8 secretion at both 3 and 6 h, mirroring its pro-apoptotic behaviour and consistent with the pro-inflammatory consequences reported for integrin-clustering antibodies such as the anti-α5β1 antibody JBS5 [7].

### JB1a restores IL-1β-disrupted ATP oscillation

Intracellular ATP levels in untreated chondrocytes showed a characteristic sinusoidal oscillation. Addition of IL-1β (10 ng/ml) reduced ATP concentration over extended time courses and abrogated this oscillation at earlier time points (Fig 2C). When IL-1β was added together with JB1a, the oscillation of ATP levels was restored, although the mean ATP concentration remained reduced. The functional significance of the residual reduction in mean ATP is not yet established.

### Ear-Punch Pilot

To explore sustained local delivery, JB1a was formulated in a polyethylene-glycol (PEG) emulsion and compared with control. Histological sections of joints from animals receiving the JB1a/PEG-emulsion depot are shown in S1 Fig, supporting the feasibility of local, sustained antibody delivery to the joint.

## Discussion

The principal finding of this study is that a single, chondrocyte-directed intervention—allosteric modulation of β1 integrin with the antibody JB1a—produces coordinated functional, structural, cellular and biochemical recovery in inflammatory arthritis. Intra-articular JB1a reversed joint swelling and hyperalgesia, restored joint architecture, reduced chondrocyte apoptosis in vivo, and in vitro abolished IL-1β-induced caspase activation, attenuated IL-8 secretion and restored the oscillatory dynamics of intracellular ATP. Together these observations position β1-integrin conformational state as an upstream determinant of chondrocyte survival and of the inflammatory tone of the joint.

A central and, in our view, mechanistically informative result is the divergence between two adhesion-blocking antibodies. JB1a and 6S6 both interfere with β1-integrin-mediated adhesion, yet they exert opposite effects on chondrocyte survival and on IL-8 secretion. This dissociation argues strongly that the therapeutic benefit of JB1a is not a consequence of adhesion blockade per se, but of the specific conformational state that it imposes on the receptor. 6S6 promotes integrin clustering and homotypic aggregation and, like the anti-α5β1 antibody JBS5, is pro-inflammatory [7]; both behave as though they drive the outside-in signalling that ECM fragmentation would be expected to trigger. JB1a does the opposite.

The molecular basis for this difference lies in where and how JB1a engages the receptor. JB1a recognises an epitope centred on residues 82–87 of the hybrid domain, and possibly residues 179–184 of β1 integrin. Integrin activation proceeds through a large-scale conformational transition in which the ectodomain extends and the hybrid domain swings out from the βI domain, an obligatory step that couples ligand binding to high-affinity signalling and to cytoskeletal engagement [14,15,16]. By binding across the hybrid domain, JB1a is expected to restrain this swing-out and thereby to stabilise the receptor in its physiological, low-affinity conformation, opposing the final steps of integrin activation [17]. Consistent with this interpretation, we have previously shown by Förster resonance energy transfer that JB1a induces an intermediate, partially extended conformation and inhibits the full activation elicited by Mn^2+^, behaving pharmacologically as an allosteric dual agonist/antagonist rather than as a conventional blocking antibody [9]. Because the resulting conformation preserves ECM binding while altering both association and dissociation kinetics, JB1a is best described as a partial antagonist that resets, rather than abolishes, β1-integrin signalling.

How does conformational restraint of β1 integrin translate into chondrocyte survival? Our data implicate the actin cytoskeleton and cellular energetics as the proximal effectors. β1-integrin-dependent adhesion organises F-actin, and the actin cytoskeleton is itself a recognised regulator of mitochondrial function and apoptosis, with ATP availability both shaping and being shaped by actin dynamics [18]. We propose that, upon local injury, degradation of the ECM promotes aggregation of cell-surface receptors including β1 integrin; this micro- or macro-aggregation initiates outside-in signalling and cytoskeletal engagement that increases F-actin assembly, raises cell-surface tethering forces and diverts ATP, changes that are known to compromise mitochondrial integrity and to sensitise cells to death. The restoration of ATP oscillation by JB1a, together with abolition of caspase 3/7 activation, is consistent with relief of this mechanical and metabolic strain. This mechanistic picture parallels our findings in a distinct organ system, where allosteric β1-integrin modulation inhibited injury-induced caspase activation, F-actin aggregation and ATP dysregulation and induced tissue repair [9,19], suggesting a conserved mechanosensing pathway that operates across tissues; consistent with this, allosteric β1-integrin modulation attenuated lesion-induced motor asymmetry in the unilateral 6-hydroxydopamine mouse model of Parkinson’s disease, albeit as proof-of-concept behavioural evidence [20].

These results also reframe how chondrocyte apoptosis relates to joint inflammation. Chondrocyte death is a feature of both osteoarthritic and inflammatory cartilage, and accumulation of apoptotic bodies that are inefficiently cleared can amplify local inflammation [21]. IL-1β is a principal catabolic cytokine acting on the chondrocyte and a driver of matrix degradation and inflammatory mediator release [22]. By interrupting the IL-1β-to-apoptosis axis at the level of integrin mechanosignalling, JB1a acts upstream of the effector programmes that current therapies address, and does so within the chondrocyte itself.

The therapeutic implications are twofold. First, JB1a defines a target space that is orthogonal to the immunomodulatory and cytokine-directed agents that dominate RA management: it is chondroprotective rather than immunosuppressive, and it acts locally. Second, the requirement for a specific receptor conformation, rather than for adhesion blockade, has direct consequences for drug design—activating or clustering ligands may be counter-productive, whereas conformation-selective, hybrid-domain-directed modulators may be therapeutically favourable. The parallel between the sinusoidal decline in hyperalgesia and the histological resolution of disease further suggests that functional read-outs may track structural repair in this setting, although the reduction in hyperalgesia should not be attributed solely to analgesia given the concomitant resolution of joint pathology.

Several limitations temper these conclusions. The histological and hyperalgesia scores are semi-quantitative and were obtained in a single, adjuvant-driven model that reproduces many but not all features of human RA; confirmation in complementary models (for example collagen-induced arthritis) and with quantitative imaging is warranted. Group sizes were modest, and the mechanistic account of the ATP–actin–mitochondrial axis, while internally consistent and supported by our prior biophysical data, remains partly inferential in cartilage. The in vitro work used primary human chondrocytes whereas the in vivo work used mice, and cross-species differences in integrin biology cannot be excluded. Finally, the residual reduction in mean ATP in JB1a-treated, IL-1β-exposed cells is unexplained and merits dedicated study. Future work should map the 6S6 epitope, define the JB1a-stabilised conformation structurally, establish dose– response and durability of the local depot formulation, and test whether allosteric β1-integrin modulation is chondroprotective in models of osteoarthritis as well as inflammatory arthritis.

In conclusion, we identify β1 integrin as a pivotal regulator of chondrocyte energy homeostasis and survival and show that allosteric modulation of the receptor through its hybrid domain reverses functional and structural injury in experimental arthritis. Taken together with our accompanying work in lung [9,19] and in a neuronal Parkinson’s disease model [20], these findings support a unifying view of integrin mechanosensing in cellular injury and identify allosteric β1-integrin modulation as a novel, mechanistically distinct therapeutic strategy for joint disease.

## Abbreviations

ANOVA: analysis of variance
ATP: adenosine triphosphate
DMARD: disease-modifying anti-rheumatic drug
DMEM/F12: Dulbecco’s modified Eagle’s medium/nutrient mixture F-12
ECM: extracellular matrix
EDTA: ethylenediaminetetraacetic acid
FCA: Freund’s complete adjuvant
FCS: fetal calf serum
FRET: Förster resonance energy transfer
H&E: haematoxylin and eosin
IgG: immunoglobulin G
IL-1β: interleukin-1β
IL-8: interleukin-8
PEG: polyethylene glycol
PI3K: phosphoinositide 3-kinase
RA: rheumatoid arthritis
TUNEL: terminal deoxynucleotidyl transferase dUTP nick-end labelling

## Acknowledgments

The authors thank the late Professor Robert J. Naylor for scientific and editorial guidance, and Professor J. A. Wilkins (University of Manitoba) for providing the JB1a antibody. Generative AI (Claude, Anthropic) was used to assist with language editing, reformatting of the manuscript to journal style, and the assembly of candidate bibliographic references; it was not used for scientific input, data generation, data analysis or interpretation. All scientific content and conclusions are those of the authors, and all references were checked against primary sources by the authors.

## Author contributions

R. AlJamal-Naylor and D. J. Harrison are co-inventors of the intellectual property describing the use and mechanism of β1 integrin in tissue repair and contributed equally to the intellectual property, scientific concept, planning and execution of the project. N. Barton and D. S. McQueen contributed to the in vivo experiments. S. McIntyre contributed to the histology, tissue culture and cytokine measurements. All authors reviewed and approved the manuscript.

## Figures

**S1 Fig.**
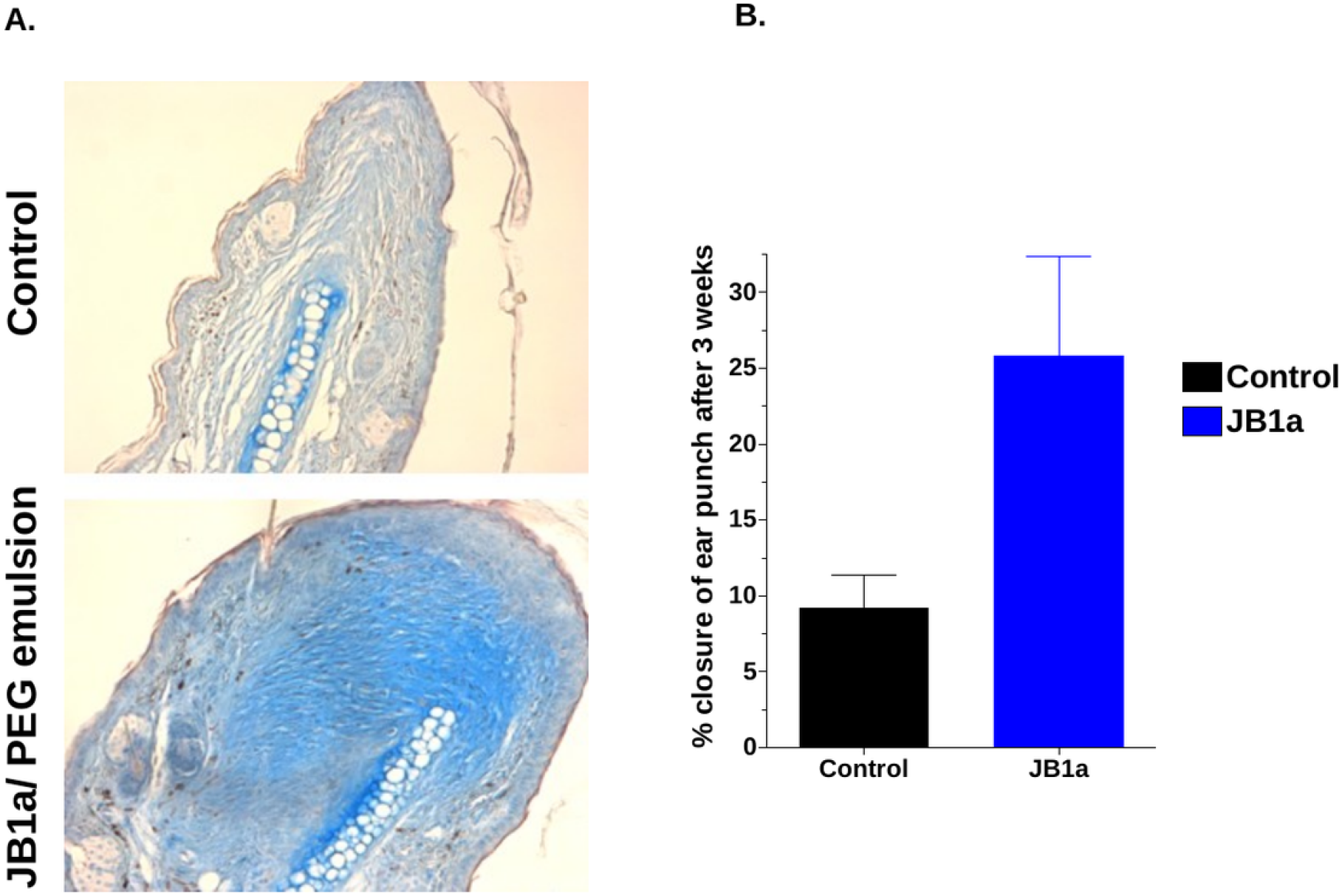
JB1a in a wound-healing / regeneration model. (A) Representative Alcian blue-stained sections of control versus JB1a/polyethylene-glycol (PEG)-emulsion-treated tissue. (B) Percentage closure of an ear-punch wound after 3 weeks in control versus JB1a-treated animals, indicating enhanced closure with JB1a.

